# Wild raccoons are more innovative but not bolder than another ecological generalist, the Virginia opossum, on a pull-string task

**DOI:** 10.1101/2025.04.21.649439

**Authors:** Kristy A. Adaway, Matthew H. Snider, Candace Vinciguerra, Caitlin A. Kupferman, James C. Beasley, F. Blake Morton

## Abstract

Humans are altering natural environments at unprecedented rates. Ecological generalism is one of the strongest predictors of survival in light of these changes which, in animals, may be shaped by bold and innovative behaviours. Species with greater habitat generalism are proposed to behave more boldly (e.g., touching and ingesting novel food). Species with greater dietary generalism are proposed to behave more innovatively (e.g., novel problem-solving to access food). Support for both hypotheses exists but remains largely restricted to broad comparisons between generalists and specialists. Further comparative data are needed to understand the extent to which these behavioural patterns might shape more nuanced ecological variation between species, such as species with finer-scale niche differences. We compared bold and innovative behaviour in two wild sympatric generalists, the Northern raccoon (*Procyon lotor*) and Virginia opossum (*Didelphis virginiana*), using a pull-string task with strings and wires attached to vertical cups containing high-value food. Although both species are broadly classified as generalists, raccoons show relatively greater habitat and dietary flexibility than opossums. Because they live sympatrically, it offers a unique opportunity to test – at the same time and locations – whether these finer-scale niche differences are reflective of differences in their bold and innovative behaviour. We predicted that raccoons would display bolder behaviour (in terms of touching our novel task) and more innovative behaviour (in terms of solving it). We found that both species were equally likely to acknowledge and touch the task, but raccoons were more likely to display innovation to access the food. Raccoons’ propensity for using innovation to solve novel foraging challenges may contribute to their greater ecological flexibility compared to opossums. More broadly, our findings may help explain differences in how each species adapts to environmental changes.

**Highlights:**

- Bold and innovative behaviour may help some species adapt to environmental changes
- We administered a novel pull-string task to wild raccoons and opossums
- Raccoons displayed more innovative, not bolder, behaviour than opossums
- Both species are generalists, but raccoons are more flexible than opossums
- Differences in innovation may shape their finer-scale niche differences

## Introduction

As a result of human activities, global biodiversity is changing at unprecedented rates (Pereira, et al., 2012). Five major types of human-induced environmental changes have been linked to this on-going “biodiversity crisis”, including habitat disturbance, pollution, overexploitation, the introduction of invasive species, and climate change (Sih, et al., 2011; Pereira, et al., 2012; Prakash & Verma, 2022). By 2050, it is estimated that 68% of the world’s human population will be urban (United Nations, 2019), which will undoubtedly lead to further habitat loss and fragmentation. In terms of biodiversity, some animals thrive in light of these on-going environmental changes, particularly invasive species and urban exploiters (Lowry, et al., 2013; Barrett, et al., 2019). Other animals, by contrast, are less resilient and either avoid environmental changes or decline in abundance whilst trying to adapt to them (Pimm, et al., 2014). To date, it is still unclear why some species are able to thrive and persist in light of on-going environmental changes; such uncertainty remains one of the most fundamental challenges to being able to predict important conservation outcomes, such as a species’ likelihood of extinction (Malhi, et al., 2020).

The initial response of an animal to changes within its environment is often behavioural (Tuomainen & Candolin, 2011). Thus, predicting a species’ behaviour plays a pivotal role in determining the extent to which a species is able to cope with environmental changes (Sih, et al., 2011; Sih, 2013; Wong & Candolin, 2015). Many of these changes will inevitably expose wildlife to situations that are temporally, physically and/or spatially novel to them. Such situations can, in many instances, offer a species with new or modified opportunities to acquire important resources such as food. Novel food opportunities can range anywhere from something relatively minor, such as discovering a new or modified resource in an unexpected location (Morton, et al., 2023), to something relatively major, such as an unexpected shift in large-scale food and habitat availability (e.g., urbanisation and climate change) (Bateman & Fleming, 2012). Exploiting novel, and hence relatively unfamiliar, food sources can expose an animal to unknown and potentially life-threatening risks (e.g., predators, traps, or toxins) (McMahon, et al., 2014; Prasher, et al., 2019). Thus, how a species responds to novel food-related opportunities, by either exploiting or avoiding them, can influence outcomes related to the survival of a species, including health, reproduction, and lifespan (Sol, et al., 2011; Oro, et al., 2013; Johnson-Ulrich, et al., 2019).

Bold behaviour, defined here in terms of an animal’s willingness to engage with unfamiliar and therefore potentially risky situations (Bergvall et al., 2011; Breck et al., 2019), may be a key factor in shaping whether or not an animal will exploit novel food opportunities. Being relatively bolder, for example, and therefore willing to approach and touch a new or modified resource, can allow some animals to take advantage of an opportunity before it is taken by a competitor (Webster et al., 2009).

Another behaviour used by animals to exploit novel food opportunities is innovation, defined here in terms of using a new or modified behaviour to solve a new or existing challenge (Lee, 1991; Reader & Laland, 2003). Having a greater tendency to innovate can improve foraging efficiency (Lee, 2003) and provide animals with greater opportunities to exploit resources that less-innovative species are unable to exploit, such as extractive foraging challenges (Reader & Laland, 2003).

There are two prevailing hypotheses to explain the occurrence of bold and innovative behaviour in relation to a species’ likelihood of exploiting novel food-related opportunities, which are based on a species’ placement along a generalism-specialism gradient. First, in terms of habitat generalism, species that are relatively bold, such as approaching and touching novel stimuli, are expected to occupy a larger range of habitats compared to other species because their behaviour allows them to be opportunistic and exploit novel food-related opportunities (Ducatez et al., 2015). Second, in terms of dietary generalism, species that are relatively innovative are expected to display greater flexibility compared to other species in terms of the types of items they exploit because innovative behaviours help them physically acquire novel resources, such as encased foods (Ducatez et al., 2015).

Ecological generalism is one of the strongest predictors of extinction, with more generalist species being at lower risk (Ducatez, et al., 2014; Senior, et al., 2021; Carballo-Morales, et al., 2024). Accounting for differences in bold and innovative behaviour, such as including these characteristics as covariates within predictive models, may therefore help improve researchers’ ability to forecast and monitor how a species might adapt to current and future environmental changes. In birds, having a greater propensity to touch and ingest unfamiliar food items (bold behaviour) and display new or modified foraging techniques (innovative behaviour) is positively associated with a species’ likelihood of colonising novel environments (Sol & Lefebvre, 2000; Sol, et al., 2002; Sol, et al., 2005). Species that are more innovative can also show greater resilience to disturbances within their environment and possess lower IUCN Red List scores (Ducatez et al., 2020). Nevertheless, other studies find that such relationships are not always consistent across species or contexts for reasons that are still unclear (Greenberg, 1990; Overington, et al., 2011a; Ducatez, et al., 2015; Henke-von der Malsburg & Fichtel, 2018; Ibáñez de Aldecoa, et al., 2024). Thus, there remains a pressing need for further studies to better understand when, how, and why bold and innovative behaviours interact with ecological generalism to shape species’ persistence.

The current study investigated the bold and innovative foraging behaviour of two wild, free-ranging North American generalists, the Northern raccoon (*Procyon lotor*) and Virginia opossum (*Didelphis virginiana*). Raccoons and opossums are interesting models to compare for a variety of reasons: First, they are sympatric species, making it possible to facilitate behavioural comparisons by experimentally giving them the same novel food-related opportunities within the same timeframe and environment. Second, evidence increasingly shows that innovative behaviours may play an important role in raccoons’ ability to thrive and persist within changing environments, such as dynamic urban settings (Daniels, et al., 2019; Stanton, et al., 2021; Stanton, et al., 2022; Stanton, et al., 2024). Third, data on the relationship between bold and innovative behaviour, particularly with regards to opportunistic foraging, are largely focused on birds and primates (Sol & Lefebvre, 2000; Sol et al., 2002; Sol et al., 2005; Ducatez et al., 2020), with fewer studies on wild mammalian carnivores (like raccoons) and marsupials (like opossums). Finally, most studies examining the relationship between ecological generalism and bold/innovative behaviours have focused largely on broad comparisons between generalist versus specialist species. Although raccoons and opossums are broadly classified as ecological generalists, both species still differ in the degree and mechanisms by which they achieve ecological breadth (Gardner & Sunquist, 2003; Gehrt, 2003). Because they live sympatrically, comparisons between raccoons and opossums offer a unique opportunity to compare what behavioural strategies might potentially shape these finer-scale niche differences.

Throughout their native range, the breadth of habitats inhabited by raccoons is far greater than that of opossums, particularly in Canada and parts of the United States (Gardner & Sunquist, 2003; Gehrt, 2003). In terms of diet, raccoons exploit some dietary items that opossums are unable or find difficult to access, such as certain types of shellfish and waterfowl (Gardner & Sunquist, 2003; Gehrt, 2003). Raccoons, but not opossums, are also renowned for acquiring items using complex extractive foraging skills, such as outdoor rubbish bins with closed lids (MacDonald & Ritvo, 2016). When similar research methods are used to compare the dietary habits of both species, raccoons can display greater dietary flexibility, with some studies reporting greater dietary divergence between urban and rural populations of raccoons but not opossums (Nicholson & Cove, 2022; Glebskiy, et al., 2024), and other studies showing greater uptake of unfamiliar food items in raccoons compared to opossums from the same locations (Helton et al., 2023; Hill et al., 2024). Finally, although comparative data on innovation in wild opossums are lacking, raccoons have relatively larger brains than opossums (Gittleman, 1986; Weisbecker & Goswami, 2010), which is an important predictor of innovation in different taxonomic groups (Benson-Amram, et al., 2016). Collectively, given these habitat and dietary characteristics, we predicted that raccoons would display bolder, more innovative behaviour compared to opossums when given an opportunity to exploit an unfamiliar food source.

## Methods

### Ethical Note

This study was ethically approved by the Animal Welfare Ethical Review Body of the University of Hull (FHS 209) and was carried out in accordance with the ASAB/ABS guidelines. No animals were handled, all trail cameras were placed away from footpaths to minimize public disturbance, and food items used to attract our study species were not harmful if ingested by other animals.

### Study Sites and Subjects

We conducted this research at a total of 135 locations throughout North and South Carolina, USA (Figure S1). Sixty-one of these locations were on the Savannah River Site, an 803 km^2^ property managed by the US Department of Energy. Fifty-one locations were in the Croatan National Forest. Fourteen locations were within and around the William B. Umstead State Park, and nine locations were on privately owned land between the coastal and piedmont regions of North Carolina. These locations were characterised by a variety of habitats found within the core geographic range of both of our study species, such as pine and hardwood forests, swamps, recreational parks, and farmland.

We used the following criteria for including locations in this study: (1) landowner permission, (2) accessibility to our study species (e.g., no barriers/fences), (3) ability to place equipment out of public view to avoid theft or vandalism, and (4) the location could not be <1 km from another study area. Because our study animals were not marked for identification, this latter criterion was used to reduce the chances of the same individuals discovering our task across multiple locations.

Since our study species were free-ranging and their participation was voluntary, some animals might have avoided our testing locations. However, our goal was to test their likelihood of being bold and innovative enough to exploit the objects, which required them to first acknowledge (at a distance) the objects before deciding whether to avoid or approach and physically interact with them. Hence, as with other studies of wild, free-ranging animals (e.g., Breck et al., 2019; Morton, 2021; Morton, et al., 2023), we based our analyses on animals that were at least able and willing to visit our study locations and walk within view of the camera’s 120° sensing angle.

## Task apparatus

We administered a modified version of a classic vertical pull-string task (Jacobs & Osvath, 2015) to each of our 135 study locations between February 2020 and November 2021. Previous research has shown that raccoons are capable of solving vertical pull-string tasks (Michels, et al., 1961). To our knowledge, Virginia opossums have not been tested with pull-string tasks prior to our study. However, opossums are capable of handling a wide variety of objects because of their dextrous paws, which is facilitated by an opposable thumb for grasping and pulling items closer to them (Krause & Krause, 2006), and indeed, studies show they can solve a range of other physical problem-solving tasks (James, 1984). In terms of size, raccoons are on average larger and taller than opossums. Despite these differences, opossums are not believed to be at a disadvantage regarding a pull-string task. For instance, the mere fact that opossums can support their own body weight whilst climbing up trees illustrates that they are strong enough to raise the very lightweight cups of the task used in the current study. While indeed the height of raccoons is larger than opossums, it is typical of opossums to use their prehensile tails to perform a range of activities, including for balance, hanging upside down from branches, and for carrying objects such as nesting materials (McManus, 1970). Thus, in our situation, the tail of opossums could be used, for example, to position themselves closer to the apparatus by hanging from the tree branches, making manipulation of the task easier, or to manipulate the apparatus and bring it closer to the branch by using the tail to hook and move along the tree branch.

The task was administered at each location for 16.75 ± 6.69 days before it was removed, regardless of whether our study species visited the task. At thirty-six of these locations, we returned 118.58 ± 81.43 days later to deploy the task again; we did this because neither of our study species, raccoons or opossums, were detected during the first sampling period. Two locations where neither raccoons nor opossums were detected in the first or second instance were resampled a third time (131.66 ± 102.61 days after the initial deployment).

The task was constructed from basic household materials, including plastic cups, strings, and plastic fishing wire. There was a total of four conditions (Figure 1). Condition 1 was a classic pull-string design, comprised of a normal white-cotton string attached to a cup that did not contain food; this allowed us to assess whether animals would attempt to interact with our task materials regardless of whether there was visual confirmation of food inside the cup. Condition 2 was comprised entirely of fishing wire attached to a cup containing food. Condition 3 was comprised of 50% string and 50% plastic fishing wire attached to a cup containing food, with the wire representing a “gap” in the centre of the string. Condition 4 was the same design as the Condition 1, comprised of a normal string attached to a cup, but this condition contained food.

**Figure 1.**
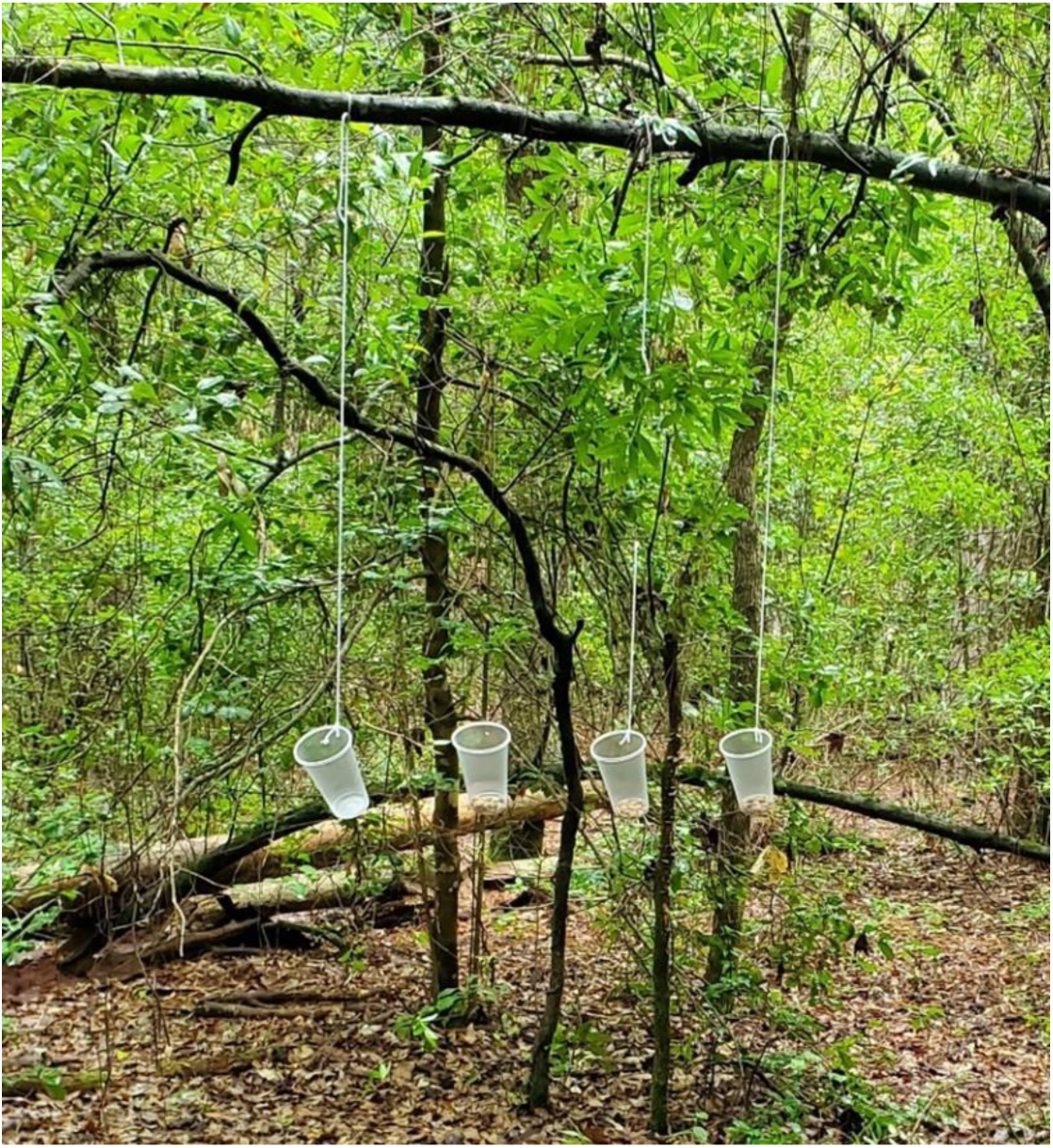
Pull-string task administered to raccoons and opossums, depicting from left to right: Condition 1 (string attached to cup, no food), Condition 2 (wire attached to cup with food), Condition 3 (string/wire combination attached to cup with food) and Condition 4 (string attached to cup with food).

The string materials used in Conditions 1, 3, and 4 were more visible to detect compared to the plastic wire materials used in Conditions 2 and 3. The wires were also more difficult for animals to handle due to the material being slippery in their coarse paws (Video S1). Thus, each of the four conditions provided participants with different options of varying difficulty to choose from, forcing them to find different ways of exploiting each condition. For example, to exploit the most difficult condition (Condition 2), animals could potentially grasp the wire with their teeth, or – for opossums – use their prehensile tail as a hook, then move away from the apparatus, bringing the cup closer to them.

The cups used for Conditions 2-4 contained either 30 dog food pellets (for *N* = 111 locations) or 30 grains of corn (for *N* = 24 locations). The two different food conditions were selected to address potential differences in motivation that may arise because of one food condition being more desirable. We selected these food items because both types are considered high-value rewards for both of our generalist species (Corn: Gardner, & Sunquist, 2003; Gehrt, 2003; Dog food: Johnson-Delaney, 2014; McWilliams & Wilson, 2015). Food was not replenished to ensure minimal human disturbance of the area. The cup in Condition 1 did not contain any food, and was used to establish whether animals would pull on the strings regardless of whether food was inside.

Raccoons and opossums are both arboreal, frequently climbing into trees to forage from branches and vines (Gardner, & Sunquist, 2003; Gehrt, 2003). Therefore, each condition was hung from an overhanging tree branch with a diameter large enough to support the weight of both species. The order in which cups were vertically hung from left to right on the tree branch was randomised across locations. The distance between each cup and the ground was approximately 110.26 ± 21.02 cm. The distance between each cup and the branch from which it was hanging was approximately 82.94 ± 18.53 cm. The distance between each of the cups was approximately 25 cm. An icepick was used to puncture a small hole into the bottom of each cup to allow rainwater to drain through. To help attract our study species to the vicinity of the puzzle, approximately 45mL of “fruit-scent” lure or fish oil was placed on a neighbouring tree branch (no more than 1.5 m away from the cups).

### Recording behaviour from trail cameras

Once the task was deployed, researchers did not return to the location until the task was scheduled to be retrieved; thus, the contents of the videos recorded at each location were unknown until testing had ended. Researchers were not present at the time of animal testing. To record animal activity, an Apeman IR trail camera (20MP) or Enkeeo PH760 infrared motion-sensor trail camera was placed adjacently on a neighbouring tree trunk approximately 3.58 ± 0.90 m away to record the subjects’ behavioural responses to the task. The trail cameras had a 120° sensing angle and a triggering distance of 20m. Our video lengths were set to record for up to 10 minutes with five second trigger delays and five second intervals between videos to ensure they would detect movement before the subjects began to interact with the task (i.e., when climbing the tree to investigate the task). Camera lenses were also sprayed with defogger and, where necessary, vegetation was removed from around the testing area to improve visibility.

### Measuring species differences in bold and innovative behaviour

Many factors can underpin the extent of bold and innovative behaviour exhibited by animals, which are not necessarily due to a single specific reason. Animals, for example, may not display such behaviour to exploit novel foraging opportunities if they are too afraid, not hungry enough, or do not persist in exploring the task long enough to discover how it operates. Importantly, however, the purpose of our study was to determine *whether* (not why) our study species would display bold and innovative behaviour, and so the only way they could do this was by physically engaging with the task and displaying those behaviours. Also, in the current study, we were not interested in trait boldness, which describes stable behavioural differences between individuals across a wide variety of contexts (Bergvall et al., 2011). Instead, as in other studies (Breck et al., 2019; Morton, 2021; Morton et al., 2023), we operationalised bold behaviour within our specific food-related context if animals physically touched the task by grabbing, pushing, pulling, and/or biting them, or making physical contact with their nose while smelling them. This is because our research question was related to whether a species was bold enough to exploit an unfamiliar food-related situation, which required them to physically touch the objects.

Since these were wild, free-ranging animals that were not tagged for reliable individual recognition, we could not control the number and frequency of visits from individuals to the puzzles. Thus, to reduce pseudo-replicates, we quantified bold and innovative behaviour using a classic method of binary coding, similar to previous studies of wild animal behaviour with unmarked individuals (e.g., Breck et al., 2019; Morton 2021; Morton et al. 2023), by scoring the presence or absence of behaviours display by each species at any point during the period in which the task was deployed at a given location. In other words, we tested, regardless of how many visits or individuals, whether a given species was more or less likely to display a certain behaviour (compared to the other species) based on the number of locations in which the behaviour was observed. This method accounts for data gaps, such as camera malfunctions or poor visibility due to fallen branches or foggy camera lenses, therefore facilitating comparability across locations. While one-zero sampling does not capture detailed continuous measures (e.g., total number of visits per species used to measure individual level traits or their underlying causes), it provides a robust method for identifying broad population-level trends in behaviour, which aligns with the goals of this study, in which we compared the likelihood of observing a given behaviour across locations (i.e., population level trends) (Martin & Bateson 2007).

To determine whether raccoons were more likely to display bolder behaviour compared to opossums, we followed previous studies of bold animal behaviour (Breck et al., 2019; Morton, 2021; Morton, et al., 2023) and calculated the likelihood of a species touching our task at any point during the period in which it was deployed at a given location. As previously discussed, here we are testing the likelihood of observing this behaviour at a given location, and not on the frequency of behaviours at a location. This was done for animals that clearly acknowledged the task by coding whether (1) or not (0) they were observed touching it at any point during the sampling period (i.e., one data point per location for each of the four task conditions). Alternative ways of measuring bold behaviour exist, such as walking speed or the latency to approach an object to within a certain body length, but as mentioned before, our research question centred on behaviours displayed at the species level, whereby populations from a given location had to physically touch the objects in order to exploit them, regardless of how long (or how many individuals) it might have taken before such behaviour was observed.

Animals were scored as having acknowledged the task if they looked directly at any of the cups or corresponding strings from any location (e.g., looking up from the ground below, looking down from the branches above, or looking across from neighbouring vegetation). They were scored as having touched the task if they physically explored any part of the task (cups, string, or wire). After conducting extensive searches across all of our study locations, we were unable to find configurations of objects that resembled the task used in our study (i.e., plastic cups hanging from trees with string and wire); meaning, animals that encountered our pull-string task were unlikely to have seen this particular set-up before. Thus, while there are, of course, other ways to characterise the “novelty” of an object, we considered our task to be novel in terms of its specific design and the locations in which animals encountered them.

For locations where animals touched the task, we scored whether (1) or not (0) they were able to ultimately gain access to the contents of a cup and noted the particular behaviour(s) used to achieve this. In terms of Condition 1 (string, no food), animals were scored as having exploited this condition if they retrieved the cup and handled it in a way that would allow them to access the food if it was available. As discussed, there were a number of ways animals might conceivably exploit the task. However, these were wild animals, and so we were unable to control how they interacted with it, leading to some behaviours being ambiguous with regards to whether or not they could be classified as “innovative”, such as an animal that might be large or tall enough to stretch its body to grab a cup without having to touch the string or wire holding it.

As discussed, our study defined innovation following Reader & Laland (2003) and Lee (1991), who describe it as using a new or modified behaviour to solve a new or existing challenge. Based on this definition, we therefore classified string- and wire-pulling as an innovation because it required subjects to perform a novel action within this unfamiliar context to retrieve food (i.e., operating the strings/wires of the task). This distinguishes string/wire pulling from pre-existing, well-rehearsed motor patterns such as simply reaching or grasping for the cup directly. While indeed additional behaviours could be used to exploit tasks that may involve a certain degree of innovation (e.g., raccoons can hang upside down to grab food), we ran separate analyses – one test with all of the behaviours that ultimately led to an animal obtaining food from the task, and one test for string/wire pulling behaviour specifically – because it was the clearest example of innovation for this task. This classification aligns with prior research in comparative animal behaviour, where string-pulling tasks are commonly used to assess innovation (Jacobs & Osvath, 2015; Alem, et al., 2016; Logan, 2016).

### Statistical analyses

To determine the inter-observer, and within-observer reliability agreement for each behaviour (i.e., acknowledge, touch, and exploit) for each of our study species, Cohen’s kappa tests were conducted. For all behaviours in each species, there was excellent inter-observer agreement (k > 0.75) between K. A., who coded all of the videos, and an independent coder who coded over 25% of videos (Tables S1-S4). Similarly, there was excellent within-observer agreement (k > 0.75) between K. A.’s original codings and over 50% of the videos re-coded over a month later (Tables S5-S8).

Cohen’s kappa tests were used to determine the inter-observer reliability agreement for which of the four cup conditions were exploited at each location. There was excellent inter-observer agreement (k > 0.75) between K.A., who coded all of the videos, and an independent coder who coded 100% of the videos where animals successfully exploited at least one of the cup conditions in the task (Table S9).

To test for significant differences in our study species’ behaviour (i.e., acknowledging, touching, or exploiting the task), we conducted Pearson’s Chi-squared tests with Yates’ continuity corrections and Fisher’s exact tests. We structured our analysis by comparing the following behaviours for each species:

- **Acknowledging the task:** we compared the number of raccoons and opossums that acknowledged the task versus those that did not, using the total number of each species that visited the area.
- **Touching the task:** we compared the number of raccoons and opossums that touched the task versus those that did not, using only the animals that had acknowledged the task.
- **Exploiting the task:** we compared the number of raccoons and opossums that exploited the task versus those that did not, using the animals that had touched the task.

We conducted Pearson’s Chi-squared tests with Yates’ continuity corrections and Fisher’s Exact tests, whilst applying the Benjamini-Hochberg procedure to account for multiple hypothesis testing to test for significant differences in raccoons’ behaviour for each of the four task conditions (Figure 1). We conducted Fisher’s exact tests to test for differences in behaviour between locations where the two different food conditions were used (i.e., dog food and corn). A Mann-Whitney U test was used to determine if there was a significant difference between species in the amount of time they spent operating the task during their first visit, i.e., when all of the cup conditions in the task were still fresh and undisturbed by other animals. Mann-Whitney U tests were also used to determine whether there was a significant difference in the length of time (in seconds) which raccoons or opossums spent at the task (measured by the length of time they spent on screen); this allowed us to evaluate the effect of species differences in sampling effort, and hence the probability of observing bold and innovative behaviour. All analyses were conducted using R, version 4.3.3 (R Core Team, 2024), and the irr package (Gamer & Lemon, 2019). All graphs were created using R, version 4.3.3 (R Core Team, 2024), and the ggplot2 (Wickham, 2016), dplyr (Wickham et al., 2023a), tidyr (Wickham, et al., 2024) and the scales packages (Wickham, et al., 2023b).

## Results

### Raccoon behavioural responses towards the pull-string task

Of the 135 locations where the task was deployed, raccoons were recorded at 51 locations (37.8%) (Figure S2). Of these locations, raccoons acknowledged the task at 37 locations (72.6%) (Figure 2). Of the locations where raccoons acknowledged the task, raccoons from 29 locations (78.4%) went on to touch the task, and of the locations where raccoons touched the task, raccoons from 21 locations (72.4%) gained access to at least one of the cup conditions (Figure 2). Raccoons from 17 of these locations used behaviours that were clearly innovative as defined by Reader and Laland (2003) (Table 1) (see behaviours 1 to 8 in Video S2).

**Figure 2.**
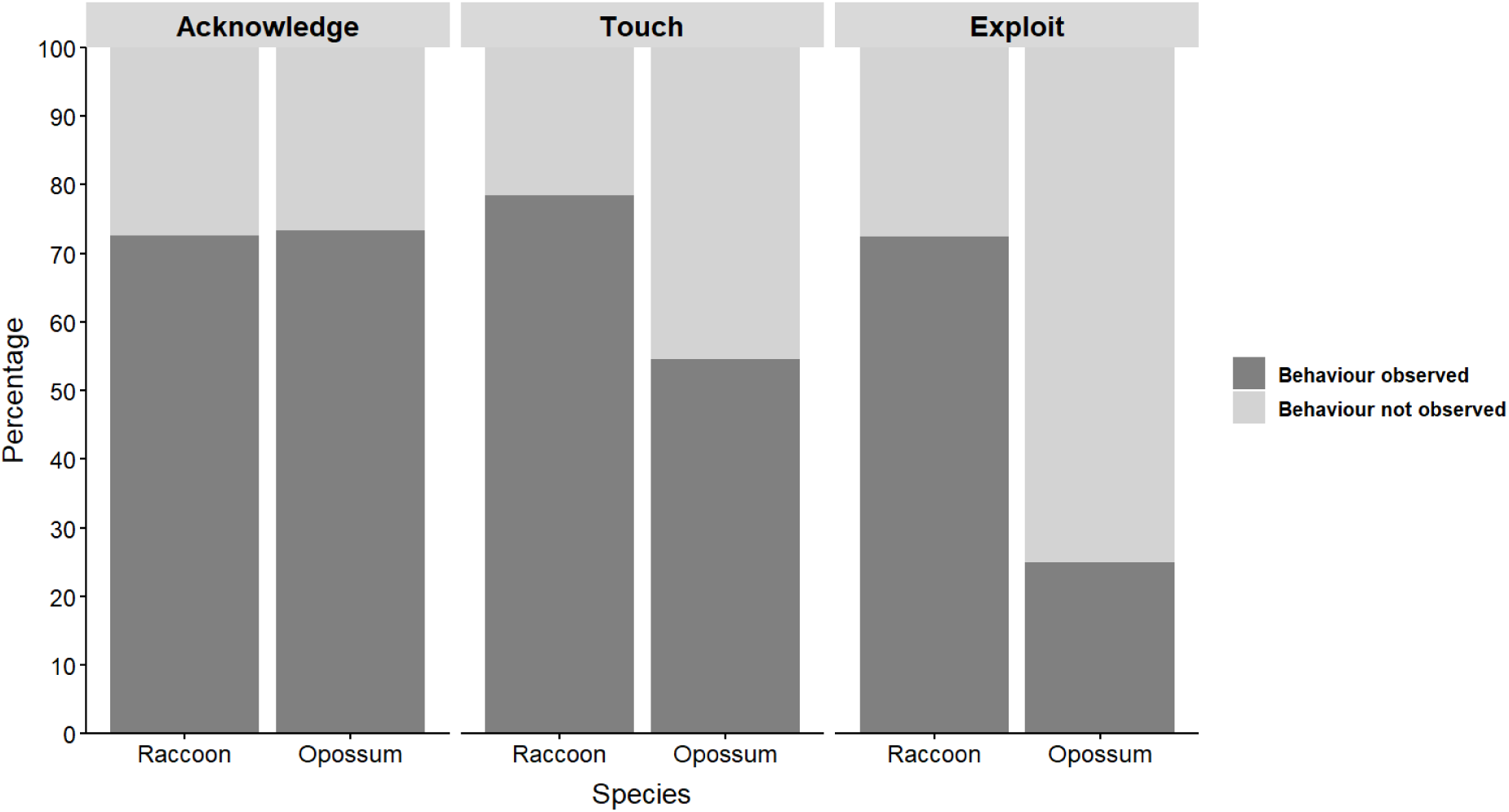
The percentage of raccoons and opossums that 1) acknowledged the task across 51 locations for raccoons and 30 locations for opossums, 2) touched the task across 37 locations for raccoons and 22 locations for opossums, and 3) exploited the task across 29 locations for raccoons and 12 locations for opossums.

**Table 1.**
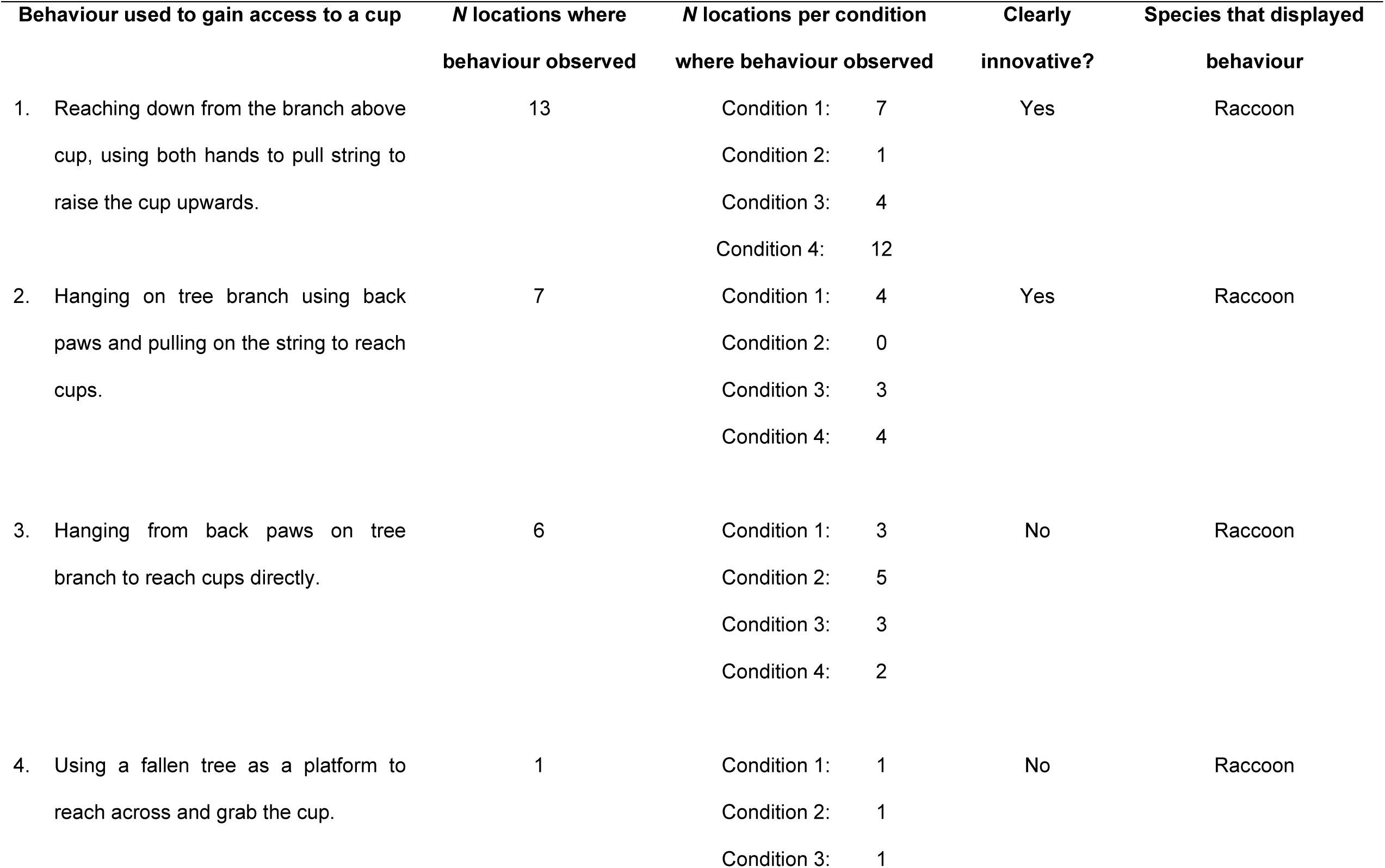
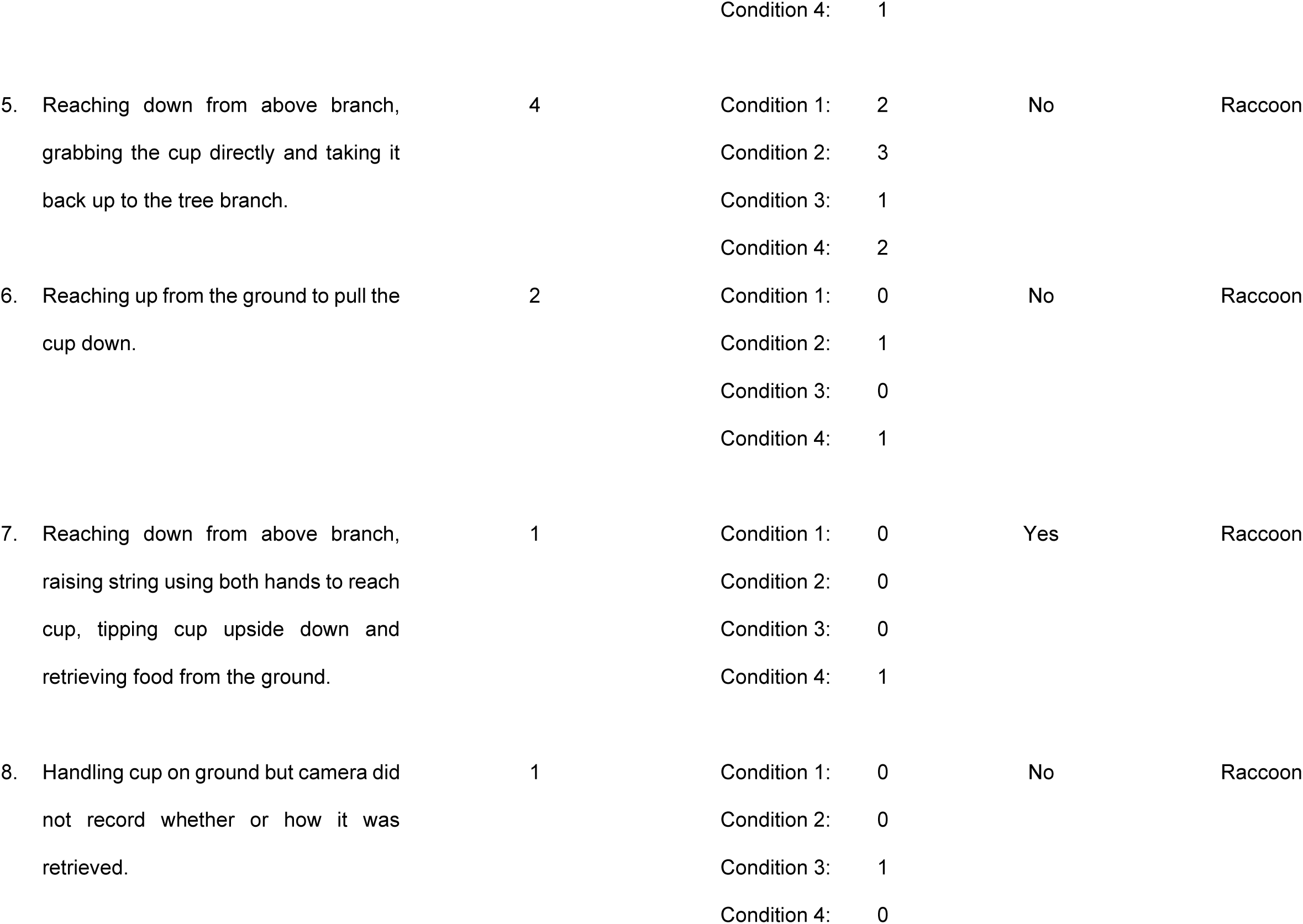
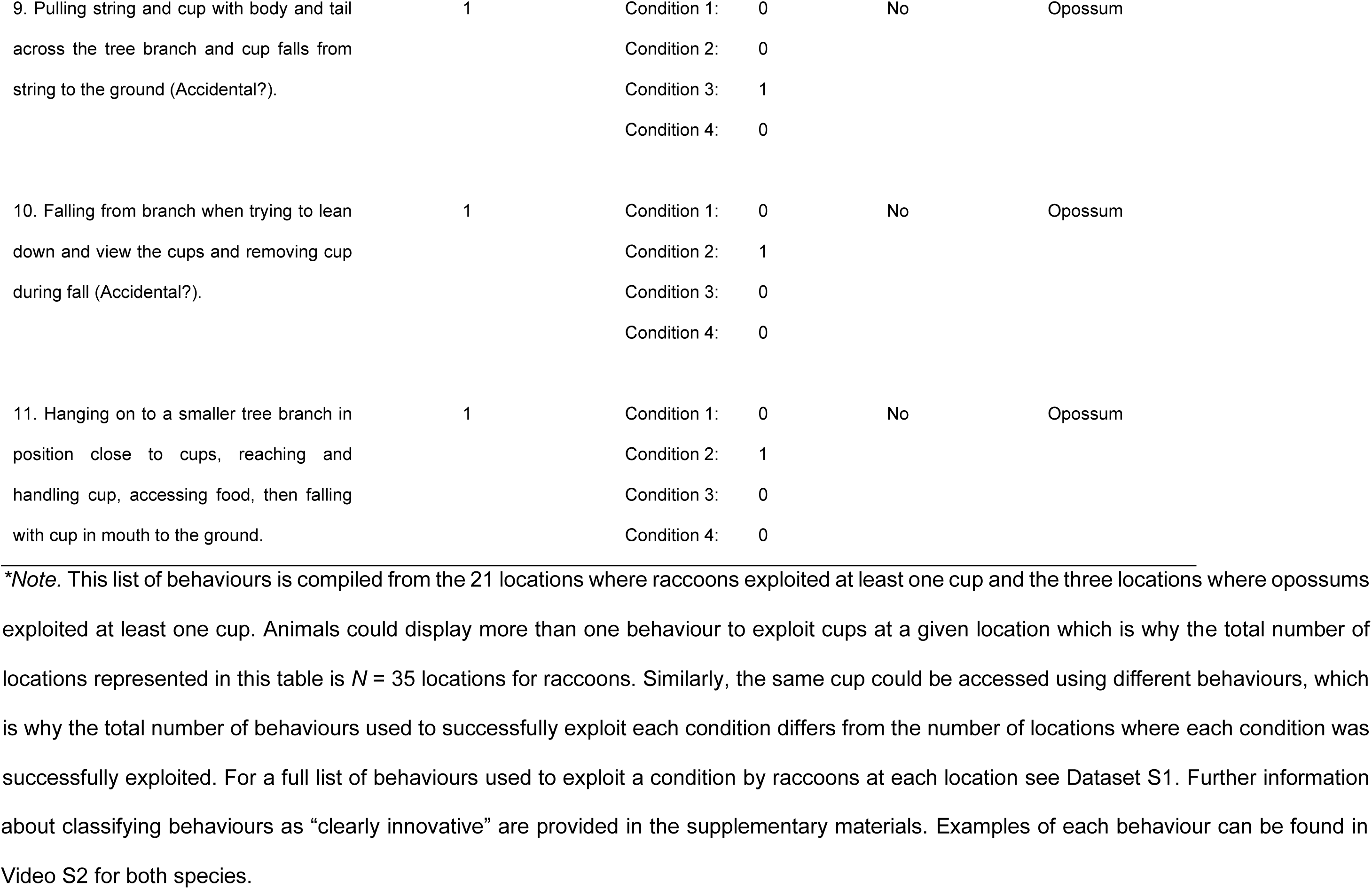
Each type of behaviour displayed by raccoons and opossums that resulted in an animal gaining access to the food rewards from at least one of the cup conditions in the task.

Raccoons interacted with the task in a range of different ways to gain access to cups (Table 1). On six occasions, two different strategies were used by raccoons to access the same cup at the same location, and both strategies led to them gaining access to some food. All of these strategies are included in Table 1, with the number of locations per condition where each behaviour was displayed.

Of the locations where raccoons successfully exploited at least one of the conditions of the task, Condition 1 (string cup with no food) (Figure 1) was exploited at 15 locations, Condition 2 (wire cup with food) was exploited at 11 locations, Condition 3 (string+wire cup with food) was exploited at 12 locations, and Condition 4 (string cup with food) was exploited at 20 locations (Figure 3). Raccoons showed a preference for exploiting Condition 4 compared to Condition 2 (Fisher’s exact test, *P* = 0.022; odds ratio: 0.065; *N* = 20 locations for Condition 2, *N* = 21 locations for Condition 4); the number of locations varied for condition 2 because on one occasion the cup was exploited by an opossum prior to the raccoon’s arrival. None of the other comparisons were significant (Table S10).

**Figure 3.**
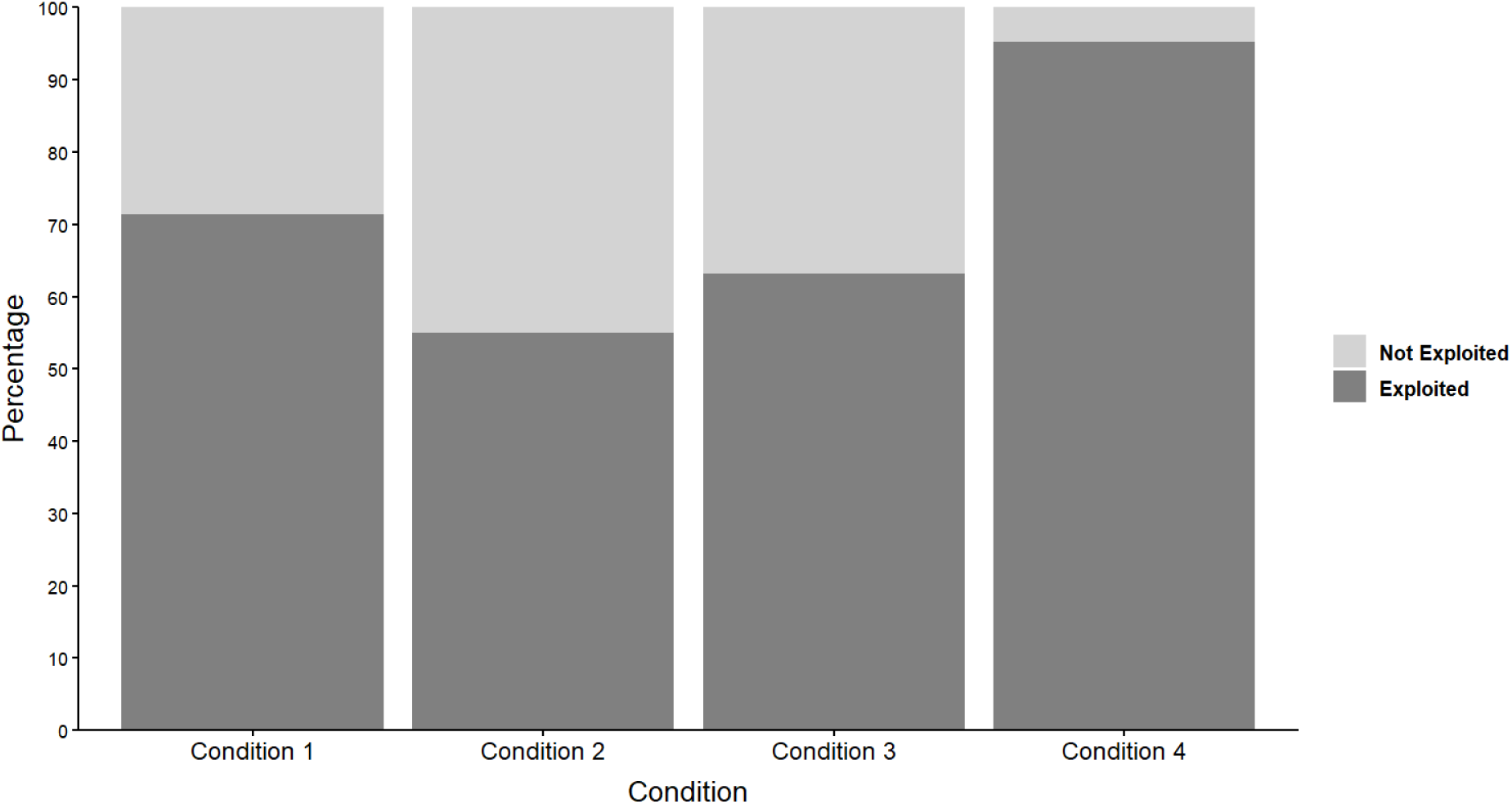
The percentage of each condition that was exploited (and not exploited) across the number of locations where raccoons successfully exploited the task. This includes data from 21 locations for Conditions 1 and 4, 20 locations for Condition 2, and from 19 locations for Condition 3. The number of locations varied per condition because, on three occasions, at least one of the cups had already been disturbed prior to the raccoon’s arrival.

We examined the order in which they exploited each condition, excluding cups that had already been disturbed or exploited by an opossum prior to the raccoon’s arrival. The order in which each condition was exploited across locations also reflects raccoons’ preference, whereby Condition 4 was exploited first in most instances and Condition 2 was exploited later, most often as the third or fourth condition (Table 2).

**Table 2.** The order and number of times each condition was exploited across a total of 21 locations where raccoons successfully exploited the task.

| <u>Condition</u> | <u>Exploited first</u> |  | <u>Exploited second</u> |  | <u>Exploited third</u> |  | <u>Exploited fourth</u> |  |
| --- | --- | --- | --- | --- | --- | --- | --- | --- |
|  | <u>N locations</u> | <u>%</u> | <u>N locations</u> | <u>%</u> | <u>N locations</u> | <u>%</u> | <u>N locations</u> | <u>%</u> |
| 1 | 5 | 23.81 | 7 | 43.75 | 3 | 21.43 | 0 | - |
| 2 | 1 | 4.76 | 0 | - | 5 | 35.71 | 5 | 71.43 |
| 3 | 5 | 23.81 | 4 | 25 | 1 | 7.14 | 2 | 28.57 |
| 4 | 10 | 47.62 | 5 | 31.25 | 5 | 35.71 | 0 | - |
| **Total locations** | **21** |  | **16** |  | **14** |  | **7** |  |

Raccoons were observed at 34 locations where dog food was used as the task reward, and at 17 locations where corn was used as the reward. Raccoons were slightly more likely to acknowledge the task if dog food was used (Fisher’s exact test: *P* = 0.045; odds ratio: 0.446; *N* = 34 locations with dog food, 17 locations with corn). However, we found no significant association between the type of reward and raccoons’ likelihood of touching (Fisher’s exact test: *P* = 0.373; odds ratio: 2.24; *N* = 28 locations with dog food, 9 locations with corn) or exploiting a cup from the task (Fisher’s exact test: *P* = 0.305; odds ratio: 3.42; *N* = 23 locations with dog food, 6 locations with corn).

### Opossum behavioural responses towards the pull-string task

Of the 135 locations where our task was deployed, opossums were detected at 30 locations (22.2%) (Figure S3), of which all but one had dog food as the task reward. Of the locations where opossums were detected, opossums from 22 locations (73.3%) acknowledged the task (Figure 2). Of these locations, opossums from 12 locations (54.6%) went on to touch the task; opossums that touched the task went on to gain access to at least one of the cups at three locations (25%) (Figure 2) (see behaviours 9 to 11 in Video S2). Of the locations where opossums successfully exploited the task, they accessed Condition 2 at two locations and Condition 3 at one location. None of the behaviours used to exploit these cups were considered to be cases of clear innovation (Table 1).

### Behavioural comparisons between species

Our species comparison revealed no significant difference between raccoons and opossums in terms of the number of locations where each species acknowledged the task (Chi-squared; *X^2^* _1_ < 0.001, *P* = 1; *N* = 51 locations with raccoons, 30 locations with opossums). Similarly, there was no significant difference between the two species in terms of the number of locations where they touched the task (Chi-squared; *X^2^* _1_ = 2.66, *P* = 0.103; *N* = 37 locations with raccoons, 22 locations with opossums). However, the first raccoon to discover the task at a given location (i.e., when all of the task conditions were fresh and undisturbed) spent on average 63.13 ± 55.38 seconds operating it before leaving, whilst the first opossum to discover the task spent on average 9.33 ± 11.66 seconds; thus, compared to opossums, raccoons were significantly more motivated to operate the task for longer periods of time compared to opossums (Mann-Whitney U test: W = 8.5, *P* = 0.005; *N* = 15 locations with raccoons, 6 locations with opossums). Raccoons that touched the task were also found to be more likely than opossums to succeed at gaining access to at least one of the cups (Fisher’s exact test: *P* = 0.013; odds ratio: 7.43; *N* = 29 locations with raccoons, 12 locations with opossums), even when only comparing the number of locations with behaviours that were clearly innovative (Table 1) (Fisher’s exact test: p < 0.001; odds ratio: Inf; *N* = 29 locations with raccoons, 12 locations with opossums).

There was no significant difference in the length of time (in seconds) raccoons or opossums spent at the task from the start to the end of the sampling period (raccoons: 360.80 seconds ± 583.62; opossums: 302.07 seconds ± 365.23) (Mann-Whitney U test: W = 735.5, *P* = 0.777, *N* = 51 locations with raccoons, 30 locations with opossums). There was also no significant difference in the length of time either species spent at the task after acknowledging its presence (raccoons: 453.08 seconds ± 644.34; opossums: 345.82 seconds ± 350.07) (Mann-Whitney U test: W = 379, *P* = 0.67, *N* = 37 locations with raccoons, 22 locations with opossums). Similarly, there was no significant difference in the length of time raccoons or opossums spent at the task between acknowledging and touching the task (raccoons: 46.55 seconds ± 68.28; opossums: 164.08 seconds ± 196.48) (Mann-Whitney U test: W = 235.5, *P* = 0.079, *N* = 29 locations with raccoons, 12 locations with opossums), or between the length of time raccoons or opossums spent at the task between acknowledging and exploiting the task (raccoons: 120.38 seconds ± 186; opossums: 122 seconds ± 162.84) (Mann-Whitney U test: W = 33, *P* = 0.93, *N* = 21 locations with raccoons, 3 locations with opossums). Some raccoons, but no opossums, acknowledged the presence of the trail camera, which was approximately 3.5m away on an adjacent tree trunk, but they did not appear to be alarmed by the camera’s presence, only briefly looking at it before returning to their main activity.

There were 13 locations where both species were present, but not at the same time. At four of these locations, neither of our study species touched or exploited the task. At four locations, an opossum visited first but did not touch or exploit the task before a raccoon visited and exploited the task, and there were no subsequent visits from opossums. At three locations, one of our study species visited and touched the task before the other study species visited the task (a raccoon at one location and an opossum at two locations); however, in these instances, the task conditions were still intact (i.e., not exploited) by the time the second species arrived, and there were no subsequent visits from the first species. There was one instance where an opossum had visited a location and exploited one condition of the task before a raccoon visited and exploited the remaining conditions.

There was also one location where opossums may have lacked opportunity to exploit the task because a raccoon visited and exploited them beforehand. Raccoons and opossums were both solitary when exploiting the task, except for three locations where two raccoons were observed visiting at the same time. At these three locations, only one of the raccoons gained access to the rewards, not both raccoons. Excluding these four locations, i.e., where an opossum lacked opportunity or conspecifics potentially influenced the other animals’ behaviour, revealed no significant change in our aforementioned species comparisons of bold and innovative behaviour (Table S11).

## Discussion

We conducted a test in the wild to determine whether, compared to opossums, raccoons were more likely to exhibit bolder behaviour (defined in terms of their likelihood of touching a novel food source) and more innovative behaviour (defined in terms of using a new or modified behaviour to exploit that source). We hypothesised that raccoons would display bolder, more innovative behaviour due to their greater habitat and dietary generalism. We found that raccoons were no more likely than opossums to acknowledge or touch the task, but they were significantly more likely to use innovation to gain access to the food rewards. Thus, in partial support of our original hypothesis, raccoons within this particular context were more likely than opossums to display innovative, but not necessarily bolder, behaviour.

Our findings for raccoons are consistent with studies using a wide variety of different methods and task designs, which show that this species is an adept problem-solver in the wild (Stanton et al., 2022; Stanton et al., 2024; Lazure & Weladji, 2024). Our findings for opossums are consistent with findings by Albert et al. (2020), which found that a sister species of the Virginia opossum, the white-eared opossum (*Didelphis albiventris*), performs similarly on vertical pull-string tasks by attempting to access the rewards directly (e.g., reaching out to get the food) rather than indirectly (e.g., pulling on the strings to bring the reward closer).

Raccoons were more likely to gain access to the cup in Condition 4 than in Condition 2. This was perhaps due to the differences in difficulty between the two conditions, which may have presented raccoons with physical and/or cognitive challenges. Condition 4, for example, consisted of a visible string attached to the cup containing a reward, whilst Condition 2 contained wire attached to the cup that was less visible and also more difficult to manipulate due to the material being too slippery on raccoons’ coarse paws, forcing raccoons to find alternative ways of exploiting this condition. Although some raccoons were able to exploit Condition 2, Morton (2021) proposed that wild raccoons may use an optimal foraging strategy favouring resources that are more easily accessible, especially when greater efforts are not needed or profitable to find food. This may offer at least one possible explanation as to why most of our raccoons were more likely to exploit Condition 4, if it was relatively easier (both physically and cognitively) for them to exploit. In support of this notion, Daniels, et al. (2019) reported that raccoons, though capable of finding multiple solutions to a novel, multi-access puzzle box, tend to solve the less challenging solutions first.

Interestingly, raccoons were no more likely to exploit any of the conditions containing food over Condition 1 (i.e., the empty cup condition). One possible explanation for this is that raccoons may not have relied primarily on visual cues when interacting with the cups. Rather, they could have associated all conditions with the presence of food based on olfactory cues alone. Raccoons have highly developed olfactory senses, which they often use to locate food (Buzuleciu, et al., 2016). Thus, it is possible that they did not visually examine the task to distinguish between food and non-food conditions before interacting with it. However, despite the significant difference between Conditions 2 and 4, there was no significant difference between Conditions 1 and 2. One possible explanation is that raccoons were considered to have exploited the condition if they retrieved the cup, and so if some raccoons were using olfactory cues to explore the task, they may have abandoned the condition (before exploiting it) once the cup was close enough to see that food was not inside. If so, then scent may have initially motivated their engagement, but raccoons may have then visually assessed reward availability as they interacted with the task.

Our study offers unique insight into the behavioural strategies that might shape how these two generalist species respond to novel environmental changes. In particular, although we found no significant difference in the likelihood of either species touching our novel foraging task, we found that raccoons were more likely to display innovative behaviour compared to opossums. Although further research is of course needed, raccoons’ propensity for using innovation to solve novel foraging challenges may contribute to their greater ecological flexibility compared to opossums.

Ingesting a novel food item requires the animal to be bold enough to touch the source of that item and, indeed, some studies find that bolder animals are more likely to approach and physically touch novel food sources compared to other animals (Frost, et al., 2007; Bergvall, et al., 2011; Morton, et al., 2023). It may be that bold behaviour is less important in shaping the niche differences of both species. Alternatively, in many landscapes where they coexist, raccoons and opossums use most habitats that are available to them. Thus, behavioural differences between these two species may exist at the fringes of their distributions where their habitat divergence is more clearly defined, such as harsh arid or cold environments (Gardner, & Sunquist, 2003; Gehrt, 2003). Finally, the relationship between ecological generalism and boldness may be specific to certain taxonomic groups or become more distinct when comparing species with very broad differences in habitat and dietary breadth (e.g. due to threshold effects and diminished returns from occupying similar niches).

### Future directions

As discussed, the current study aimed to test *whether*, not why, raccoons would display bolder and more innovative foraging behaviour compared to opossums because these outward behaviours are what ultimately may help some species adapt to environmental changes, regardless of the underlying reason. To understand the “why”, further work will be needed: For instance, both of our study species have a strong sense of smell, and so it would be interesting to incorporate a combination of controlled field tests with time series analyses to test whether scent marks leftover from prior visitations played a role in future animals’ behavioural responses to novelty.

Detailed research on our study animals’ food preferences is another avenue worth investigating in the future. Although we found no evidence that task rewards had an impact on raccoons’ behaviour within this study, raccoons were more likely to acknowledge the task when dog food was used, which may be attributed to the stronger odour of the dog food compared to corn. Both of our chosen task rewards are considered high-value rewards for both of our study species (Corn: Gardner, & Sunquist, 2003; Gehrt, 2003; Dog food: Johnson-Delaney, 2014; McWilliams & Wilson, 2015). However, because of the low sample size of opossums detected at locations where corn was used as the task reward, comparisons could not be made for the species. Thus, we encourage comparative studies on the dietary preferences of wild raccoons and opossums since nuanced differences in their willingness to consume the food rewards in our study may have contributed to our findings.

Understanding the psychological underpinnings of outwardly bold and innovative behaviour, and including multiple measures of behaviour within one’s analysis, may shed further light on its relationship with ecological generalism. In particular, we encourage further studies to build upon our findings by testing, with a larger sample size of each study species, a range of other tasks varying in their degree of novelty and complexity (Carter, et al., 2012; Beckmann & Biro, 2013; Henke-von der Malsburg & Fichtel, 2018; Ibáñez de Aldecoa, et al., 2024) since these characteristics can of course play important roles in shaping the expression of bold and innovative behaviours (Daniels, et al., 2019; Lazure & Weladji, 2024; Vincze, et al., 2024). Tasks to help tease apart the underlying reason(s) for raccoons’ greater persistence in operating the task compared to opossums would be particularly insightful, since this may be reflective of greater motivation (e.g., willingness to put effort into exploiting a task), risk-taking (e.g., willingness to spend longer touching a novel, potentially risky, food source) (Takola, et al., 2021), or indeed a combination of both. If risk-taking plays a role in raccoons’ greater persistence on tasks compared to opossums, then this would offer stronger support for the notion that raccoons display relatively bolder behaviour compared to opossums.

Interactions between bold and innovative behaviour facilitate an animals’ ability to adapt to novel environments or utilise novel resources (Reader & Laland, 2003; Overington, et al., 2011b; Morton, et al., 2023). The purpose of our study was to understand the animals’ responses to novelty, and so within this context, raccoons and opossums needed to be bold enough to interact with the task before they could exploit its contents. Other studies have found that reduced boldness can prevent animals from being innovative (Overington, et al., 2011b; Mazza & Guenther, 2021; Morton, et al., 2023), whereas other studies find that bolder animals are not necessarily more innovative (Griffin & Diquelou, 2015; Morton, et al., 2023). As with these latter studies, we found no evidence that racoons (the more innovative species) were bolder than opossums. Thus, the relationship between bold behaviour and innovative behaviour is mixed between studies and species for reasons that are unclear, which future research should explore further.

Our study supports the proposal that species differences in behavioural innovation may help explain differences in their likelihood of adapting to novel environmental changes, particularly changes that result in new or modified food-related opportunities. Nevertheless, further work is still needed to better understand when, where, and how innovation might play a role in shaping a species’ responses to novel environmental changes. Henke-von der Malsburg and Fichtel (2018), for example, found that the generalist grey mouse lemur (*Microcebus murinus*) and the more specialised Madame Berthe’s mouse lemur (*Microcebus berthae*) used behavioural innovation to solve novel foraging opportunities, but the specialist species was more efficient at finding a novel solution when the challenge was more familiar. Similarly, Ibáñez de Aldecoa, et al. (2024) used a multi-access box with four opening mechanisms to compare the innovative abilities of four species of Darwin’s finch – two diet specialists that were extractive foragers and two diet generalists that were non-extractive foragers. They found that the two species of dietary specialists were faster innovators than the two species of dietary generalists. Thus, multiple measures of innovative behaviour, ranging in complexity and degree of novelty, are likely needed to understand how innovation links to animals’ responses to novel situations. Distinguishing between the manner in which innovations are defined and quantified will also be important, as well as differences in species’ opportunities and overall “need” for innovation (Lee, 2003).

### Conclusions

Bold and innovative behaviours may help some species adapt to environmental changes. Our findings contribute to the growing literature on behavioural flexibility in wild animals by comparing the bold and innovative behaviour of two wild, sympatric generalist species. We found that raccoons behaved more innovatively than opossums when exploiting novel foraging opportunities. Although both species are classified as ecological generalists, raccoons are relatively more flexible than opossums. Differences in behavioural innovation may potentially shape these finer-scale niche differences. More broadly, our findings may help explain differences in how each species adapts to environmental changes.

## Data Availability

All data are provided in dataset S1 in the Supplementary material.

## Acknowledgments

We would like to thank the students and assistants that helped to collect data, and the landowners for giving us permission to collect data on their land, particularly the U.S. Forest Service of the U.S. Department of Agriculture. We thank K. Sutter and J. Anderson (Uni Hull) for coding videos for the inter-observer reliability tests. Special thanks go to the team from the Croatan Ranger District for their support throughout data collection within the Croatan National Forest. FBM and KA thank the University of Hull for funding. Contributions of JCB and CK were partially supported by the U.S. Department of Energy under award number DE-EM0005228 to the University of Georgia Research Foundation. We are very grateful to the two anonymous reviewers for taking time to provide all of their helpful comments.

## Supplementary Material

Supplementary material associated with this article is available in the online version.

