## Supplementary materials for "Wild raccoons are more innovative but not bolder than another ecological generalist, the Virginia opossum, on a pull-string task"

**Supplementary material (Adaway et al. 2025)**

**Video S1.** Raccoon trying to solve Condition 2 of a pull-string task. <https://www.youtube.com/watch?v=PFUiuT_YJb0&feature=youtu.be>

**Video S2.** Videos of behaviours from Table 1. <https://www.youtube.com/watch?v=De86LQggxDc>

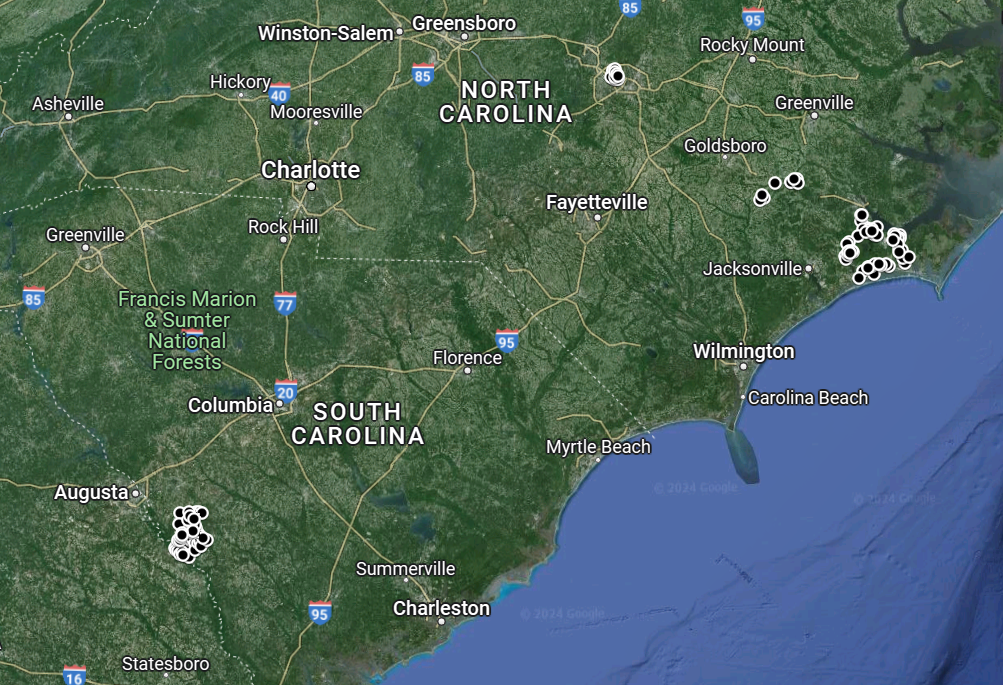

**Figure S1.** Map of the 135 research locations. Sixty-one locations at the field site of the Savanna River Ecology Laboratory. Fifty-one locations in the Croatan National Forest. Fourteen locations in the William B. Umstead State Park, and nine locations on privately owned land between the coastal and piedmont regions of North Carolina.

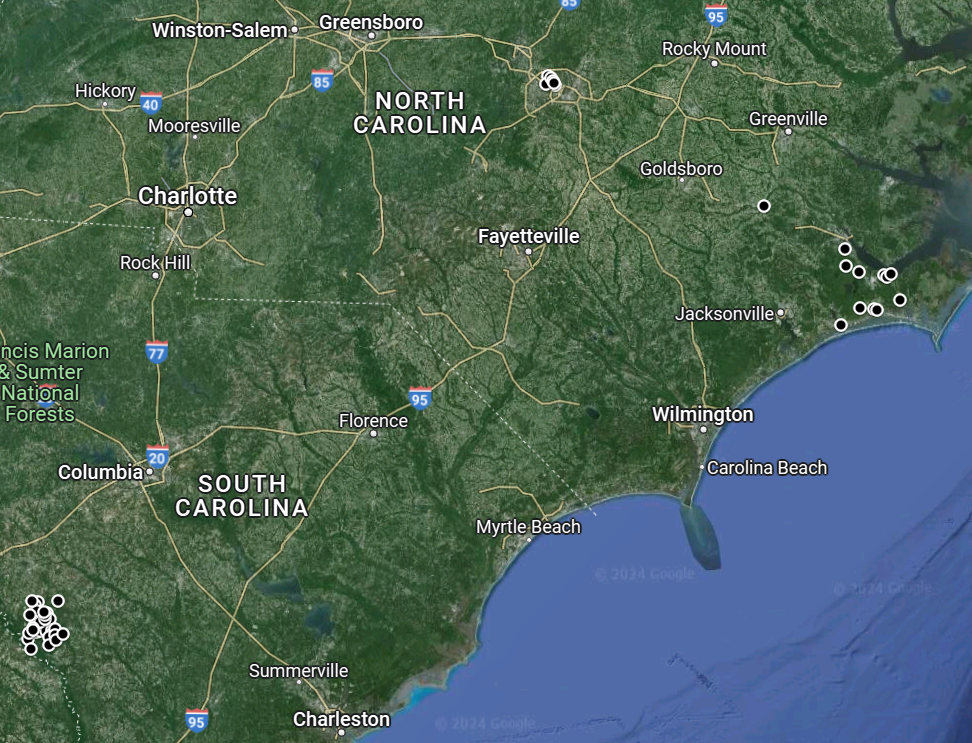

**Figure S2.** Map of the 51 locations where raccoons were present.

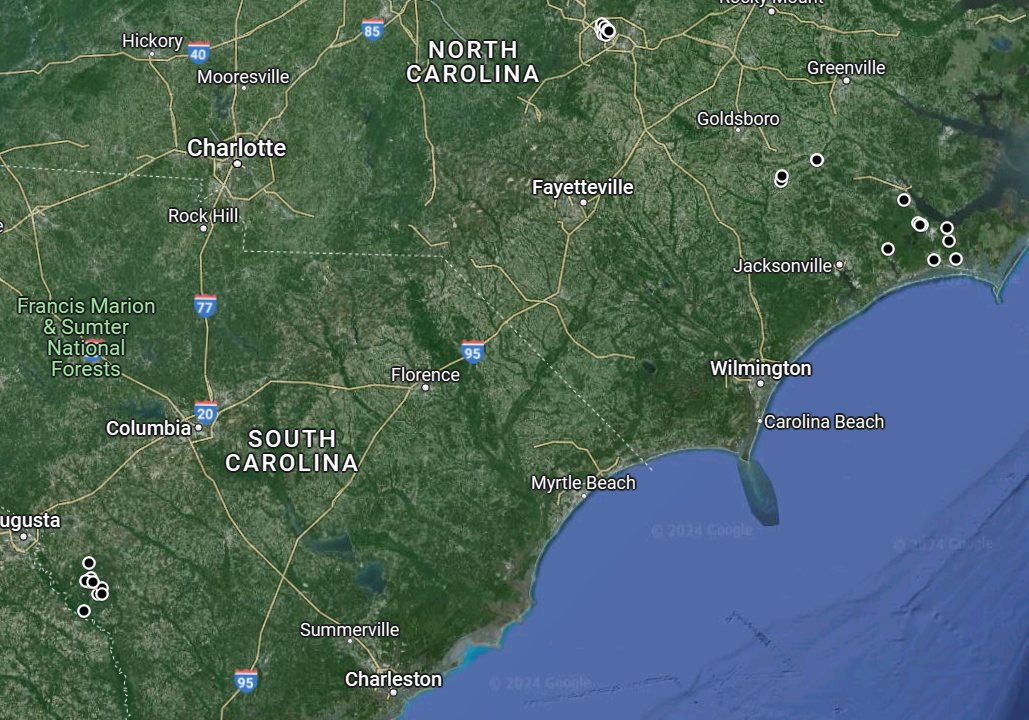

**Figure S3.** Map of the 30 locations where opossums were present.

**Table S1.** Reliability test between K.A. and an independent coder for raccoons’ acknowledgment of the task.

|  | Acknowledge task? | |
| --- | --- | --- |
| Video | Coder 1 (K.A.) | Coder 2 |
| 1 | 1 | 1 |
| 2 | 1 | 1 |
| 3 | 1 | 1 |
| 4 | 1 | 1 |
| 5 | 1 | 1 |
| 6 | 1 | 1 |
| 7 | 1 | 1 |
| 8 | 1 | 1 |
| 9 | 0 | 0 |
| 10 | 0 | 0 |
| 11 | 0 | 0 |
| 12 | 1 | 1 |
| 13 | 0 | 0 |
| 14 | 1 | 1 |
| 15 | 0 | 0 |
| 16 | 1 | 1 |
| 17 | 0 | 0 |
| 18 | 1 | 1 |
| 19 | 1 | 1 |
| 20 | 1 | 1 |
| 21 | 1 | 1 |
| 23 | 1 | 1 |
| 24 | 1 | 1 |
| 25 | 1 | 1 |
| 26 | 1 | 1 |
| 27 | 1 | 1 |
| 28 | 1 | 1 |
| 29 | 1 | 1 |
| 30 | 1 | 1 |
| 31 | 1 | 1 |

1 = behaviour observed, 0 = behaviour not observed. Interobserver reliability between both coders was K = 1.

**Table S2.** Reliability test between K.A. and an independent coder for opossums’ acknowledgment of the task.

|  | Acknowledge task? | |
| --- | --- | --- |
| Video | Coder 1 (K.A.) | Coder 2 |
| 1 | 1 | 1 |
| 2 | 1 | 1 |
| 3 | 1 | 1 |
| 4 | 1 | 1 |
| 5 | 1 | 1 |
| 6 | 1 | 1 |
| 7 | 0 | 0 |
| 8 | 0 | 0 |
| 9 | 1 | 1 |
| 10 | 1 | 1 |
| 11 | 1 | 1 |
| 12 | 1 | 1 |
| 13 | 1 | 1 |
| 14 | 1 | 1 |
| 15 | 1 | 1 |

1 = behaviour observed, 0 = behaviour not observed. Interobserver reliability between both coders was K = 1.

**Table S3.** Reliability test between K.A. and an independent coder for raccoons’ touching or exploiting the task.

|  | Touch | | Exploiting | |
| --- | --- | --- | --- | --- |
| Video | Coder 1 (K.A.) | Coder 2 | Coder 1 (K.A.) | Coder 2 |
| 1 | 1 | 1 | 1 | 1 |
| 2 | 1 | 1 | 1 | 1 |
| 3 | 1 | 1 | 0 | 0 |
| 4 | 1 | 1 | 1 | 1 |
| 5 | 1 | 1 | 1 | 1 |
| 6 | 0 | 0 | 0 | 0 |
| 7 | 0 | 0 | 0 | 0 |
| 8 | 1 | 1 | 1 | 1 |
| 9 | 1 | 1 | 0 | 1 |
| 10 | 0 | 0 | 0 | 0 |
| 11 | 0 | 0 | 0 | 0 |
| 12 | 1 | 1 | 0 | 0 |
| 13 | 1 | 1 | 1 | 1 |
| 14 | 0 | 0 | 0 | 0 |
| 15 | 0 | 0 | 0 | 0 |
| 16 | 1 | 1 | 1 | 1 |
| 17 | 1 | 1 | 1 | 1 |
| 18 | 1 | 1 | 1 | 1 |
| 19 | 1 | 1 | 1 | 1 |
| 20 | 1 | 1 | 1 | 1 |
| 21 | 1 | 1 | 1 | 1 |
| 22 | 1 | 1 | 1 | 1 |
| 23 | 1 | 1 | 0 | 1 |
| 24 | 1 | 1 | 1 | 1 |

1 = behaviour observed, 0 = behaviour not observed. Interobserver reliability between coders for raccoon touch was K = 1, interobserver reliability between coders for raccoon exploit was K = 0.824.

**Table S4.** Reliability test between K.A. and an independent coder for opossums’ touching or exploiting the task.

|  | Touch | | Exploit | |
| --- | --- | --- | --- | --- |
| Video | Coder 1 (K.A.) | Coder 2 | Coder 1 (K.A.) | Coder 2 |
| 1 | 1 | 1 | 0 | 0 |
| 2 | 1 | 1 | 0 | 0 |
| 3 | 0 | 0 | 0 | 0 |
| 4 | 0 | 0 | 0 | 0 |
| 5 | 1 | 1 | 0 | 0 |
| 6 | 0 | 0 | 0 | 0 |
| 7 | 0 | 0 | 0 | 0 |
| 8 | 1 | 1 | 0 | 0 |
| 9 | 1 | 1 | 0 | 0 |
| 10 | 0 | 0 | 0 | 0 |
| 11 | 1 | 1 | 1 | 1 |
| 12 | 0 | 1 | 0 | 0 |
| 13 | 1 | 1 | 0 | 0 |

1 = behaviour observed, 0 = behaviour not observed. Interobserver reliability between coders for opossum touch was K = 0.843, interobserver reliability between coders for opossum exploit was K = 1.

**Table S5.** Test-retest consistency for raccoons' acknowledgment of the task.

|  | Acknowledge task? | |
| --- | --- | --- |
| Video | Time 0 | Time 1 |
| 1 | 1 | 1 |
| 2 | 1 | 1 |
| 3 | 1 | 1 |
| 4 | 1 | 1 |
| 5 | 1 | 1 |
| 6 | 1 | 1 |
| 7 | 1 | 1 |
| 8 | 0 | 0 |
| 9 | 0 | 0 |
| 10 | 0 | 0 |
| 11 | 1 | 1 |
| 12 | 1 | 1 |
| 13 | 0 | 0 |
| 14 | 1 | 1 |
| 15 | 0 | 0 |
| 16 | 0 | 0 |
| 17 | 1 | 1 |
| 18 | 1 | 1 |
| 19 | 0 | 0 |
| 20 | 0 | 0 |
| 21 | 1 | 1 |
| 22 | 0 | 0 |
| 23 | 1 | 1 |
| 24 | 1 | 1 |
| 25 | 1 | 1 |
| 26 | 1 | 1 |
| 27 | 1 | 1 |
| 28 | 1 | 1 |
| 29 | 1 | 1 |
| 30 | 1 | 1 |
| 31 | 1 | 1 |
| 32 | 1 | 1 |
| 33 | 1 | 1 |
| 34 | 1 | 1 |
| 35 | 1 | 1 |
| 36 | 1 | 1 |
| 37 | 1 | 1 |
| 38 | 1 | 1 |
| 39 | 1 | 1 |
| 40 | 1 | 1 |
| 41 | 1 | 1 |
| 42 | 1 | 1 |
| 43 | 1 | 1 |
| 44 | 1 | 1 |
| 45 | 1 | 1 |

1 = behaviour observed, 0 = behaviour not observed. Agreement between K.A.'s scores at Time 0 and 1 was K = 1. Data were coded by K.A. at two different time periods separated by several months to test for intra-observer consistency.

**Table S6.** Test-retest consistency for opossums' acknowledgment of the task.

|  | Acknowledge task? | |
| --- | --- | --- |
| Video | Time 0 | Time 1 |
| 1 | 1 | 1 |
| 2 | 0 | 0 |
| 3 | 1 | 1 |
| 4 | 0 | 0 |
| 5 | 0 | 0 |
| 6 | 0 | 0 |
| 7 | 0 | 0 |
| 8 | 0 | 0 |
| 9 | 1 | 1 |
| 10 | 1 | 1 |
| 11 | 1 | 1 |
| 12 | 1 | 1 |
| 13 | 1 | 1 |
| 14 | 1 | 1 |
| 15 | 1 | 0 |
| 16 | 1 | 1 |
| 17 | 1 | 1 |
| 18 | 1 | 1 |
| 19 | 1 | 1 |
| 20 | 1 | 1 |
| 21 | 1 | 1 |
| 22 | 0 | 0 |
| 23 | 1 | 1 |
| 24 | 0 | 0 |
| 25 | 1 | 1 |
| 26 | 1 | 1 |
| 27 | 1 | 1 |
| 28 | 1 | 1 |
| 29 | 0 | 0 |

1 = behaviour observed, 0 = behaviour not observed. Agreement between K.A.'s scores at Time 0 and 1 was K = 0.922. Data were coded by K.A. at two different time periods separated by several months to test for intra-observer consistency.

**Table S7.** Test-retest consistency for raccoons’ touching or exploiting the task.

|  | Touch | | Exploit | |
| --- | --- | --- | --- | --- |
| Video | Time 0 | Time 1 | Time 0 | Time 1 |
| 1 | 1 | 1 | 0 | 0 |
| 2 | 1 | 1 | 1 | 1 |
| 3 | 1 | 1 | 0 | 0 |
| 4 | 0 | 0 | 0 | 0 |
| 5 | 0 | 0 | 0 | 0 |
| 6 | 1 | 1 | 0 | 0 |
| 7 | 1 | 1 | 1 | 1 |
| 8 | 1 | 1 | 0 | 0 |
| 9 | 0 | 0 | 0 | 0 |
| 10 | 1 | 1 | 0 | 0 |
| 11 | 0 | 0 | 0 | 0 |
| 12 | 0 | 0 | 0 | 0 |
| 13 | 0 | 0 | 0 | 0 |
| 14 | 1 | 1 | 0 | 0 |
| 15 | 1 | 1 | 1 | 1 |
| 16 | 1 | 1 | 1 | 1 |
| 17 | 0 | 0 | 0 | 0 |
| 18 | 1 | 1 | 1 | 1 |
| 19 | 0 | 0 | 0 | 0 |
| 20 | 1 | 1 | 1 | 1 |
| 21 | 1 | 1 | 1 | 1 |
| 22 | 1 | 1 | 1 | 1 |
| 23 | 1 | 1 | 1 | 1 |
| 24 | 1 | 1 | 1 | 1 |
| 25 | 1 | 1 | 1 | 1 |
| 26 | 1 | 1 | 1 | 1 |
| 27 | 1 | 1 | 1 | 1 |
| 28 | 1 | 1 | 1 | 1 |
| 29 | 1 | 1 | 0 | 1 |
| 30 | 1 | 1 | 1 | 1 |
| 31 | 1 | 1 | 1 | 1 |
| 32 | 1 | 1 | 1 | 1 |
| 33 | 1 | 1 | 1 | 1 |
| 34 | 1 | 1 | 0 | 0 |
| 35 | 1 | 1 | 1 | 1 |
| 36 | 1 | 1 | 1 | 1 |

1 = behaviour observed, 0 = behaviour not observed. Agreement between K.A.'s scores at Time 0 and 1 was K = 1 for touch, and K = 0.943 for exploit. Data were coded by K.A. at two different time periods separated by several months to test for intra-observer consistency.

**Table S8.** Test-retest consistency for opossums’ touching or exploiting the task.

|  | Touch | | Exploit | |
| --- | --- | --- | --- | --- |
| Video | Time 0 | Time 1 | Time 0 | Time 1 |
| 1 | 1 | 1 | 0 | 0 |
| 2 | 0 | 0 | 0 | 0 |
| 3 | 0 | 0 | 0 | 0 |
| 4 | 1 | 1 | 0 | 0 |
| 5 | 1 | 1 | 0 | 0 |
| 6 | 1 | 1 | 0 | 0 |
| 7 | 0 | 0 | 0 | 0 |
| 8 | 1 | 1 | 1 | 1 |
| 9 | 0 | 0 | 0 | 0 |
| 10 | 0 | 1 | 0 | 0 |
| 11 | 1 | 1 | 1 | 1 |
| 12 | 0 | 0 | 0 | 0 |
| 13 | 1 | 1 | 0 | 0 |
| 14 | 1 | 1 | 0 | 0 |
| 15 | 0 | 0 | 0 | 0 |
| 16 | 1 | 1 | 0 | 0 |
| 17 | 1 | 1 | 0 | 0 |
| 18 | 0 | 0 | 0 | 0 |
| 19 | 0 | 0 | 0 | 0 |

1 = behaviour observed, 0 = behaviour not observed. Agreement between K.A.'s scores at Time 0 and 1 was K = 0.894 for touch, and K = 1 for exploit. Data were coded by K.A. at two different time periods separated by several months to test for intra-observer consistency.

**Table S9.** Reliability test between K.A. and an independent coder for which of the four conditions raccoons gained access to at each of the locations where they successfully exploited the task.

| Location | Condition (1-4) | Coder 1 (K.A.) | Coder 2 |
| --- | --- | --- | --- |
| 1 | 1 | 1 | 1 |
|  | 2 | 1 | 1 |
|  | 3 | 1 | 1 |
|  | 4 | 1 | 1 |
| 2 | 1 | 1 | 1 |
|  | 2 | 1 | 1 |
|  | 3 | 0 | 0 |
|  | 4 | 1 | 1 |
| 3 | 1 | 1 | 1 |
|  | 2 | 0 | 0 |
|  | 3 | 0 | 0 |
|  | 4 | 1 | 1 |
| 4 | 1 | 1 | 1 |
|  | 2 | 1 | 1 |
|  | 3 | 1 | 1 |
|  | 4 | 1 | 1 |
| 5 | 1 | 1 | 1 |
|  | 2 | 1 | 1 |
|  | 3 | 1 | 1 |
|  | 4 | 1 | 1 |
| 6 | 1 | 0 | 0 |
|  | 2 | 0 | 0 |
|  | 3 | 1 | 1 |
|  | 4 | 1 | 0 |
| 7 | 1 | 0 | 0 |
|  | 2 | 0 | 0 |
|  | 3 | 0 | 0 |
|  | 4 | 1 | 1 |
| 8 | 1 | 1 | 1 |
|  | 2 | 1 | 1 |
|  | 3 | 1 | 1 |
|  | 4 | 1 | 1 |
| 9 | 1 | 1 | 1 |
|  | 2 | 0 | 1 |
|  | 3 | 1 | 1 |
|  | 4 | 1 | 1 |
| 10 | 1 | 1 | 1 |
|  | 2 | 0 | 0 |
|  | 3 | 1 | 1 |
|  | 4 | 1 | 1 |
| 11 | 1 | 0 | 0 |
|  | 2 | 0 | 0 |
|  | 3 | 0 | 0 |
|  | 4 | 1 | 1 |
| 12 | 1 | 1 | 1 |
|  | 2 | 1 | 1 |
|  | 3 | 0 | 0 |
|  | 4 | 1 | 1 |
| 13 | 1 | 0 | 0 |
|  | 2 | 0 | 0 |
|  | 3 | 0 | 0 |
|  | 4 | 1 | 1 |
| 14 | 1 | 0 | 0 |
|  | 2 | 0 | 0 |
|  | 3 | 0 | 0 |
|  | 4 | 1 | 1 |
| 15 | 1 | 1 | 0 |
|  | 2 | 0 | 0 |
|  | 3 | 1 | 1 |
|  | 4 | 1 | 1 |
| 16 | 1 | 1 | 1 |
|  | 2 | 1 | 1 |
|  | 3 | 1 | 1 |
|  | 4 | 1 | 1 |
| 17 | 1 | 1 | 1 |
|  | 2 | 1 | 1 |
|  | 3 | 1 | 1 |
|  | 4 | 1 | 1 |
| 18 | 1 | 1 | 1 |
|  | 2 | 1 | 1 |
|  | 3 | 0 | 0 |
|  | 4 | 1 | 1 |
| 19 | 1 | 1 | 1 |
|  | 2 | 1 | 1 |
|  | 3 | 1 | 1 |
|  | 4 | 1 | 1 |
| 20 | 1 | 0 | 0 |
|  | 2 | 0 | 0 |
|  | 3 | 0 | 0 |
|  | 4 | 1 | 1 |
| 21 | 1 | 1 | 1 |
|  | 2 | 1 | 1 |
|  | 3 | 1 | 1 |
|  | 4 | 0 | 0 |

1 = behaviour observed, 0 = behaviour not observed. Interobserver reliability between both coders was K = 0.917

**Table S10.** Matrix showing the likelihood of each task condition being exploited at a given location at any point during the period in which the task was available. All p-values have been adjusted by applying the Benjamini-Hochberg procedure to account for multiple hypothesis testing.

| Condition | 2 | 3 | 4 |
| --- | --- | --- | --- |
| 1 | Pearson’s chi-squared:  *X^2^* _1_ = 0.589,  *P* = 0.664 | Pearson’s chi-squared:  *X^2^* _1_ = 0.048,  *P* = 0.848 | Fisher’s exact test:  Odds ratio: 0.13,  *P* = 0.186 |
| 2 |  | Pearson’s chi-squared:  *X^2^* _1_ = 0.037,  *P* = 0.848 | Fisher’s exact test:  Odds ratio: 0.065,  ***P* = 0.022** |
| 3 |  |  | Fisher’s exact test:  Odds ratio: 0.091,  *P* = 0.052 |

*Note*: Of the 21 locations where raccoons exploited the task, we were able to obtain data on each the four conditions they exploited. This included data from 21 locations for Conditions 1 and 4, data from 20 locations for Condition 2, and data from 19 locations for Condition 3. The number of locations varied per condition because on three occasions at least one of the cups had already been disturbed or exploited by an opossum prior to the raccoon’s arrival.

**Table S11.** Comparison of the likelihood of raccoons and opossums acknowledging, touching and exploiting the task when excluding four locations: three locations where raccoons were not solitary upon exploiting the task and one location where a raccoon exploited the task before an opossum had acknowledged the task.

| Behaviour comparison | Chi-squared results | Interpretation |
| --- | --- | --- |
| Acknowledging | *X^2^* _1_ < 0.001, *P* = 1; *N* = 47 locations (raccoons), 28 locations (opossums) | Raccoons no more likely to acknowledge the task than opossums |
| Touching | *X^2^* _1_ = 0.81473, *P* = 0.367; *N* = 33 locations (raccoons), 20 locations (opossums) | Raccoons no more likely to touch the task than opossums |
| Exploiting | *X^2^* _1_ = 4.429, ***P* = 0.035**; *N* = 25 locations (raccoons), 12 locations (opossums) | Raccoons more likely to exploit the task than opossums |

**Further information about classifying behaviours as “clearly innovative”:**

In total, 11 different types of behaviour were observed being used by raccoons or opossums to gain access to at least one of the task conditions (Table 1). For 9 of these behaviours, there was 100% agreement between the original coder (KA) and three independent coders in terms of which ones were, and were not, cases of clear innovation as defined by Reader & Laland (2003).

For the remaining two behaviours (Behaviours 9 and 11 in Table 1), some of the independent coders scored them as cases of clear innovation (one coder for Behaviour 9 and two coders for Behaviour 11, respectively), whereas the other coders did not. After some discussion, it was agreed amongst all coders that these two behaviours were too ambiguous to be classified as innovative because the underlying reason for the animals’ behaviour was unclear. For example, for Behaviour 9, the opossum’s tail may have accidentally become tangled, causing the cup to fall. For Behaviour 11, the opossum may have accidentally fallen from the branch, causing the cup to fall with it.
